# A scalable pipeline for local ancestry inference using tens of thousands of reference haplotypes

**DOI:** 10.1101/2021.01.19.427308

**Authors:** Eric Y. Durand, Chuong B. Do, Peter R. Wilton, Joanna L. Mountain, Adam Auton, G. David Poznik, J. Michael Macpherson

## Abstract

Ancestry deconvolution is the task of identifying the ancestral origins of chromosomal segments of admixed individuals. It has important applications, from mapping disease genes to identifying loci potentially under natural selection. However, most existing methods are limited to a small number of ancestral populations and are unsuitable for large-scale applications.

In this article, we describe Ancestry Composition, a modular pipeline for accurate and efficient ancestry deconvolution. In the first stage, a string-kernel support-vector-machines classifier assigns provisional ancestry labels to short statistically phased genomic segments. In the second stage, an autoregressive pair hidden Markov model corrects phasing errors, smooths local ancestry estimates, and computes confidence scores.

Using publicly available datasets and more than 12,000 individuals from the customer database of the personal genetics company, 23andMe, Inc., we have constructed a reference panel containing more than 14,000 unrelated individuals of unadmixed ancestry. We used principal components analysis (PCA) and uniform manifold approximation and projection (UMAP) to identify genetic clusters and define 45 distinct reference populations upon which to train our method. In cross-validation experiments, Ancestry Composition achieves high precision and recall.

## 1 Introduction

An individual’s genome can be viewed as a mosaic of chromosomal segments of potentially different ancestries (Falush et al., 2003; Tang et al., 2005). Ancestry deconvolution, or local ancestry inference (LAI), is the task of resolving the ancestral origin of such segments. Robust ancestry deconvolution enables several important lines of research, including admixture-based mapping of disease genes (Seldin et al., 2011); disease-association studies, in which controlling for population structure is essential (Price et al., 2010); and studies of population history (Novembre and Ramachandran, 2011; Hellenthal et al., 2014).

As LAI is a classification problem, there are two general approaches one may take. One approach is generative, in which one models the joint probability of haplotypes and ancestries, and the other is discriminative, in which one models the conditional probability of ancestries given haplotype data (see section 4 for a fuller discussion of this distinction). Several ancestry-deconvolution methods implement generative models in which the ancestry of chromosomal segments is inferred using hidden Markov models (e.g., Tang et al., 2006; Price et al., 2009). Other generative approaches have used sliding-window algorithms (e.g. Pasaniuc et al., 2009). In contrast to generative approaches, discriminative approaches do not attempt to fully model the underlying admixture process. Instead, they attempt to learn directly from segments of known ancestry the conditional distribution of ancestries given haplotype data. Discriminative models make fewer assumptions about the demographic process underlying admixture and typically scale better to large datasets (Omberg et al., 2012; Kumar et al., 2020). A number of discriminative approaches have been described (Brisbin et al., 2012; Omberg et al., 2012; Maples et al., 2013; Kumar et al., 2020; Montserrat et al., 2020).

Previous LAI methods have also differed in how they treat chromosome phase. Some early methods relied on unphased genotype data and predicted for each genetic marker whether zero, one, or two alleles derive from a specified ancestral population (e.g., Pasaniuc et al., 2009; Sundquist et al., 2008; Price et al., 2009). However, there is much information to be gleaned from phase. An individual may inherit very different ancestries from their mother and father, and the ability to represent and model these differences provides greater power for identifying ancestries and enables parental contributions to be distinguished. Some methods incorporate phase information by prephasing genotypes (Brisbin et al., 2012; Maples et al., 2013). Others phase the input genotypes as part of the analysis (Bercovici et al., 2012). However, even with the availability of population-scale genetic datasets (e.g., Bycroft et al., 2018; McCarthy et al., 2016) and breakthroughs in statistical phasing methodology (e.g., Loh et al., 2016), it remains challenging to recover chromosome-scale phase information. Thus, it is beneficial to model phase errors as part of the data generation process.

In this article, we present an update to *Ancestry Composition*, a modular two-stage LAI pipeline originally described in Durand et al. (2014). In a manner similar in spirit to (Omberg et al., 2012; Maples et al., 2013), Ancestry Composition uses a discriminative approach to determine the ancestral origins of short chromosomal segments. It subsequently corrects these assignments with a generative model that jointly models the true ancestries of each haplotype, correlations in local assignments, and errors in haplotype phase inference. We have trained Ancestry-Composition models using a reference panel of more than 14,000 individuals with known ancestry, most of whom are customers of 23andMe, Inc., a personal genomics company. Ancestry Composition achieves high precision and recall when labeling chromosomal segments from more than 45 worldwide populations. In contrast, most existing LAI methods tend to be limited to a few ancestral populations and typically lack power to distinguish between closely related populations (Pasaniuc et al., 2009). We designed the method to function in a online setting in which an analyst or consumer product must continuously predict new individuals. As such, Ancestry Composition scales linearly with the number of individuals to analyze.

## 2 Methods

Ancestry Composition consists of two largely independent modules:

1. A local classifier module, wherein a string-kernel support-vector-machines classifier (SKSVM) assigns provisional ancestry labels to short locally phased genomic regions.
2. An error-correction module, wherein an autoregressive pair hidden Markov model (APHMM) corrects phasing errors, smooths provisional ancestry estimates, and assigns confidence scores.

### 2.1 Local classifier

The task of the local classifier is to assign each marker along each haplotype to one of *K* reference populations. The local classifier starts by splitting each haplotype into *S* windows of *M* biallelic markers. Each window is treated independently and is assumed to have a single ancestral origin. Thus, for each haplotype, the local classifier returns a vector *c*_1:*S*_, where *c*_*i*_ ∈ {1 …*K*} is the hard-clustering value assigned to window *i*. We implemented the local classifier using string-kernel support vector machines, a discriminative classifier.

#### 2.1.1 String-kernel support vector machine

Support Vector Machines (SVMs) are a class of supervised learning algorithms first introduced by Vapnik (1998). In its most basic form, an SVM is a non-probabilistic binary linear classifier. That is, it learns a linear decision boundary that can be used to discriminate between two classes. SVMs can be extended to problems that are not linearly separable using the soft-margin technique. For more details, see Cristianini and Shawe-Taylor (2000).

Consider a set of training data {(x_*i*_, *y*_*i*_)}_1:*N*_, where for each *i*, x_*i*_ is a feature vector in ℝ^*d*^ and *y*_*i*_ ∈ {−1, 1} is a class label. The SVM learns the decision boundary by solving the following quadratic programming optimization problem:

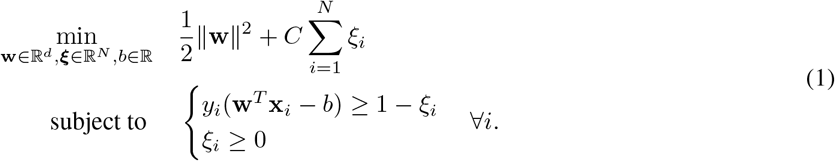

*C* is a tuning parameter that, in practice, we generally set to 1.

##### Encoding the feature vectors

In our application, each feature vector x_*i*_ represents the encoding of a haplotype window of *M* biallelic markers from a prephased haplotype. One natural encoding is to use one feature per marker, with each feature encoding the presence or absence of the minor allele. However, this encoding fails to capture the the spatial relationship of consecutive markers within the window (i.e., the linkage pattern), a distinguishing feature of genetic variation. Instead, we use every possible *k*-mer (*k* ∈ {1 …*M*}) as our features. For a haplotype segment of *M* biallelic markers, there are 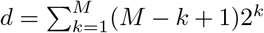 possible *k*-mers. When *M* = 100, *d* is on the order of 10^30^, so it is not feasible to directly construct feature vectors with this many dimensions. In the next section, we introduce a string kernel that enables working with our high-dimensional feature set.

##### String kernel

A key property of SVMs is that solving (1) is equivalent to solving the following dual quadratic programming problem:

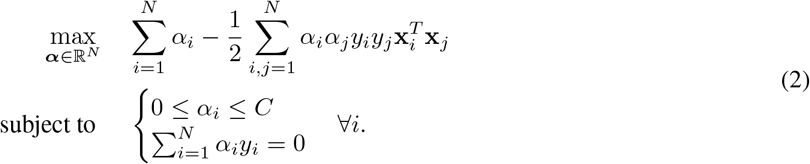

The dual representation of the SVM optimization problem depends only on the inner product 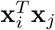, which means we can introduce kernels (Boser et al., 1992). Kernels provide a way to map observations to a high-dimensional feature space, thereby offering an enormous computational advantage, as they can be evaluated without explicitly calculating feature vectors. Denoting the input space as *χ*, let *ϕ* : *χ* → {0, 1}^*d*^ be the mapping such that for any segment *x* of length *M, ϕ*(*x*) is the vector whose elements denote the presence or absence of each of the *d* possible *k*-mers in *x*. We define our string kernel as, ∀*i, j* ∈ {1, …, *N*},

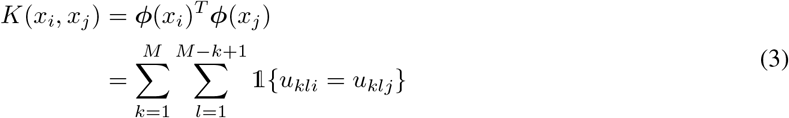

where *u*_*kli*_ is the *k*-mer starting at position *l* in haplotype window *i*. Our kernel is a special case of the weighted degree kernel (Rätsch et al., 2006). Standard dynamic programming techniques can be used to evaluate *K*(*x*_*i*_, *x*_*j*_) in *O*(*M*) operations without explicitly enumerating the *d* features for each mapped input vector. Thus, the string kernel enables us to extract a large amount of information from each haplotype window.

##### Multiclass SVMs SVMs

are fundamentally binary classifiers, but in this setting we are concerned with deciding among 45 possible populations. To assign a single hard-clustering value *k* ∈ {1, …, *K*} to a haplotype window, we trained 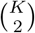 classifiers, one for each pair of populations. We assign each haplotype segment to a single population using a straightforward majority vote across all pairs. We also experimented with a one-vs-all approach that did not perform as well.

#### 2.1.2 Training data

We trained the local classifier on ~14,400 unrelated individuals, each with unadmixed ancestry from one of *K* = 45 reference populations (Table 1). This reference panel includes ~11,800 research-consented 23andMe customers, ~600 individuals from non-customer 23andMe datasets, and ~2000 individuals from publicly available datasets, including the 1000 Genomes Project (The 1000 Genomes Project Consortium, 2015) and the Human Genome Diversity Panel (Cann et al., 2002).

**Table 1:** Reference population sample composition. For Mongolia & Manchuria, the count of individuals from the 23andMe database is between one and five, with the totals in the margins reflecting a count of five in that cell.

| Population | 23andMe | Public | Total |
| --- | --- | --- | --- |
| Americas | 12 | 62 | 74 |
| Arabia | 257 | 0 | 257 |
| Ashkenazi Jewish | 1007 | 0 | 1007 |
| Balkans & Greece | 614 | 0 | 614 |
| Bengal & Northeast India | 166 | 80 | 246 |
| Biaka, Mbuti & San | 0 | 41 | 41 |
| Britain & Ireland | 935 | 79 | 1014 |
| Caucasus, Assyria & Iran | 400 | 0 | 400 |
| Central & West Europe | 936 | 21 | 957 |
| Central Asia | 113 | 24 | 137 |
| China | 544 | 251 | 795 |
| Chinese Dai | 0 | 82 | 82 |
| Congo | 597 | 0 | 597 |
| Copt | 120 | 0 | 120 |
| Cyprus | 158 | 0 | 158 |
| East Europe | 761 | 24 | 785 |
| Egypt Other | 185 | 0 | 185 |
| Ethiopia & Eritrea | 171 | 0 | 171 |
| Finland | 279 | 85 | 364 |
| Gujarat Subgroup | 85 | 67 | 152 |
| Italy | 482 | 114 | 596 |
| Japan | 346 | 128 | 474 |
| Kerala | 191 | 0 | 191 |
| Korea | 341 | 0 | 341 |
| Levant | 348 | 52 | 400 |
| Maghreb | 307 | 25 | 332 |
| Melanesia | 0 | 29 | 29 |
| Mongolia & Manchuria | $\leq 5$ | 18 | $\sim 23$ |
| Myanmar, Thailand, Cambodia & Indonesia | 69 | 8 | 77 |
| Nigeria | 54 | 226 | 280 |
| Northern India & Southern Pakistan | 254 | 46 | 300 |
| Philippines & Austronesia | 164 | 0 | 164 |
| Sardinia | 0 | 25 | 25 |
| Scandinavia | 631 | 0 | 631 |
| Senegal, The Gambia & Guinea | 23 | 135 | 158 |
| Siberia | 0 | 22 | 22 |
| Sierra Leone, Liberia, Ivory Coast & Ghana | 196 | 85 | 281 |
| Somalia | 150 | 0 | 150 |
| Southern Brahmin | 292 | 8 | 300 |
| Southern East Africa | 15 | 109 | 124 |
| Southern India Other & Sri Lanka | 204 | 85 | 289 |
| Spain & Portugal | 344 | 13 | 357 |
| Sudan | 189 | 0 | 189 |
| Turkey | 359 | 0 | 359 |
| Vietnam | 114 | 83 | 197 |
| Total | $\sim 12,418$ | 2027 | $\sim 14,445$ |

To ensure that all the reference individuals were distantly related, we used the method described in Henn et al. (2012) to estimate identity-by-descent (IBD) sharing between each pair of individuals and removed individuals from the sample until no pair shared more than an 100 cM. We then conducted principal components analysis (PCA) and uniform manifold approximation and projection (UMAP; McInnes et al., 2018; Diaz-Papkovich et al., 2019) to identify population structure, which, when paired with survey data and analyzed jointly with the well-curated external reference panels, enabled us to define our 45 reference populations and flag outliers for removal.

For most reference populations, the research-consented 23andMe customers reported in survey responses that their four grandparents were born in a single country. For regions with large multiethnic countries (e.g., South Asia), we also required that an individual’s four grandparents either spoke a single regional language or were born in one state. Free-text responses on grandparental national, ethnic, religious, or other identities enabled us to construct reference panels for populations not defined by specific geographic regions (e.g., Ashkenazi Jews).

#### 2.1.3 Window size

A key assumption of the local classifier is that the haplotype segment within each window derives from a single population. Thus, the window-size parameter influences the timing of the admixture we can address. For example, if we sought only to infer “local” admixture in first-generation admixed individuals, then windows could potentially span entire chromosomes. More generally, if we assume a simple admixture model in which two reference populations mixed *T* generations ago, then the expected length of a single-ancestry segment is 100*/*(2*T*) cM.

Phasing switch errors also limit the sizes of segments we can consider. If a switch error occurs within a haplotype window, our assumption that the haplotype segment covered by the window has a single ancestry may no longer be valid. Thus, it is necessary to choose a window size small enough to ensure that most windows are free of switch errors. On the other hand, longer windows contain more information, which increases the power of the SKSVM to separate reference populations.

For the analyses presented in this article, we used a window size of 300 markers. This corresponds to ~0.6 cM per window and, on a genotyping platform measuring ~540,000 markers, divides the genome into ~1800 windows. We chose this window size because we find it provides a good balance between retaining ancestry-related information within windows and precluding recombination events and phasing errors within them.

### 2.2 Error-correction module

The local classifier generates noisy ancestry estimates, so we developed an error-correction module to smooth hard-clustering assignments using information from adjacent windows. To compute smoothed assignment probabilities, we have implemented an autoregressive pair hidden Markov model (APHMM) that explicitly represents both haplotypes covering a genomic window. With *S* denoting the number of windows of *M* markers, consider a directed probabilistic graphical model (Figure 1) consisting of:

**Figure 1:**
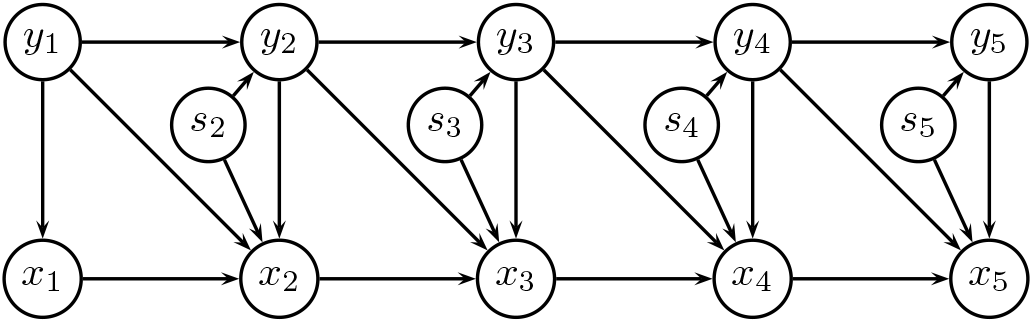
Graphical model of the error-correction module for sequence of length *S* = 5.

- *S* hidden states, *y*_1:*S*_ = (*y*_1_, *y*_2_, …, *y*_*S*_), where 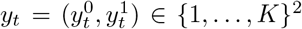 represents the true population labels of haplotypes 0 and 1 within window *t*;
- *S* observed states, *x*_1:*S*_ = (*x*_1_, *x*_2_, …, *x*_*S*_), where 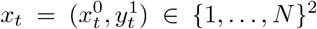 represents the observed population labels for the *t*-th pair of haplotype windows (i.e., the output from the local classifier); and
- *S* − 1 hidden switch indicators, *s*_2:*S*_ = (*s*_2_, *s*_3_, …, *s*_*S*_), where *s*_*t*_ ∈ {0, 1} denotes whether a phasing switch error has occurred between windows *t* − 1 and *t*.

Note that we implicitly assume that phasing switch errors occur only at the boundaries between windows. We model the joint probability of *y*_1:*S*_, *x*_1:*S*_, and *s*_2:*S*_ as:

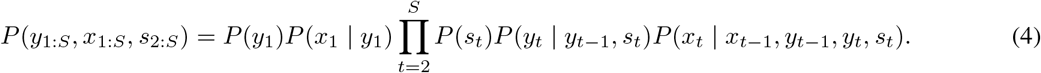

We parameterize our model with 2(*K*^2^ + *K*) + 1 parameters:

- {*µ*_*y*_}_1:*K*_, the prior distribution of hidden states following a recombination event;
- 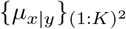, the prior distribution of emissions, conditional on hidden states;
- *σ*, the prior probability that a phasing switch error occurs between two consecutive windows;
- {*θ*_*y*_}_1:*K*_, the prior probabilities of recombination between two consecutive windows, when the first has hidden state *y*; and
- 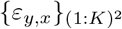, the prior probabilities of observed-state label resets between two consecutive windows, when the first has observed state *x* and both have hidden state *y*.

We express each component of the joint probability expression (4) in terms of these parameters:

1. **Initial hidden-state distribution.** We assume that the population assignments for each of the two haplotypes is sampled independently from the stationary distribution of hidden states, ***π***:

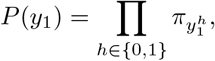

where

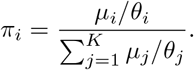 We note that, in the original version of Ancestry Composition (Durand et al., 2014), the prior probability of recombination was a scalar, *θ*, constant across hidden states. When this was the case, the stationary distribution of hidden states, *π*, was equal to the prior distribution of hidden states, ***µ***.
2. **Initial emission distribution.** The initial emissions for each haplotype are sampled independently from the prior distribution for emissions:

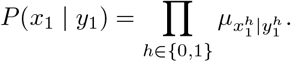
3. **Switch error model.** We assume that switch errors occur with constant probability *σ* between each pair of states:

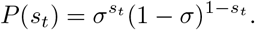
4. **Transition probability model.** For each haplotype, a recombination occurs from hidden state *y* with probability *θ*_*y*_, and for each recombination, we draw a new hidden population label from the prior distribution for hidden states. Thus,

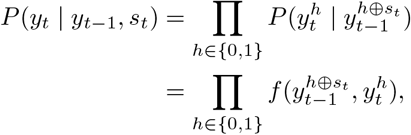

where

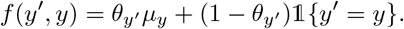
5. **Emission probability model.** In order to accommodate correlated errors in local ancestry classifications, we designed the APHMM’s emission model to be autoregressive: given no change in hidden state, the observed states are correlated. In our testing, this autoregressivity increased posterior decoding accuracy without any apparent performance decline. As with the transition probability model, we treat each haplotype independently in the emission probability model. Consider two consecutive hidden states, *y*_*t*−1_ and *y*_*t*_. If they are unequal (i.e., a true ancestry switch has occurred), then an observed-state label reset necessarily occurs, and the emission at window *t, x*_*t*_, is drawn from the prior distribution for emissions, 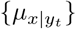. If a true ancestry switch has not occurred (i.e., *y*_*t*−1_ = *y*_*t*_ ≡ *y*), an observed-state label reset occurs with probability 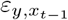:

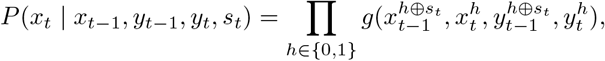

where

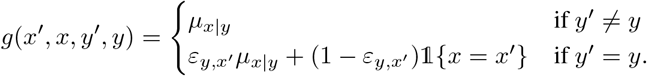

We estimate these model parameters using the expectation-maximization (EM) algorithm (Dempster et al., 1977). Posterior probabilities for each window are estimated using the forward and backward algorithms for hidden Markov models. Using dynamic programming techniques, the complexity of the posterior decoding step is *O*(*SK*^2^), where *S* is the number of windows to decode and *K* is the number of populations.

### 2.3 Posterior aggregation for hierarchical classification

Intracontinental local ancestry inference is a challenging problem, and it may not always be possible to confidently determine whether a segment derives from Scandinavia or the British Isles, either because we lack power or because the corresponding haplotypes occur at similar frequencies within the two populations. In such cases, it is often possible to confidently determine that the segment derives from a specific broader region (e.g., Northern Europe). Therefore, we have defined a four-level population hierarchy that groups populations within continents and regions (Figure 2). The *K* leaves (i.e., terminal nodes) of our hierarchy correspond to the *K* reference populations, and the highest level consists of a single root node representing the union of all populations. Broadly, the levels beneath the root correspond to continental-scale, regional-scale, and sub-regional–scale populations, respectively. Leaf nodes may occur at any of these levels; for example, Melanesia is placed immediately below the root, at the continent scale, and is not further subdivided.

**Figure 2:**
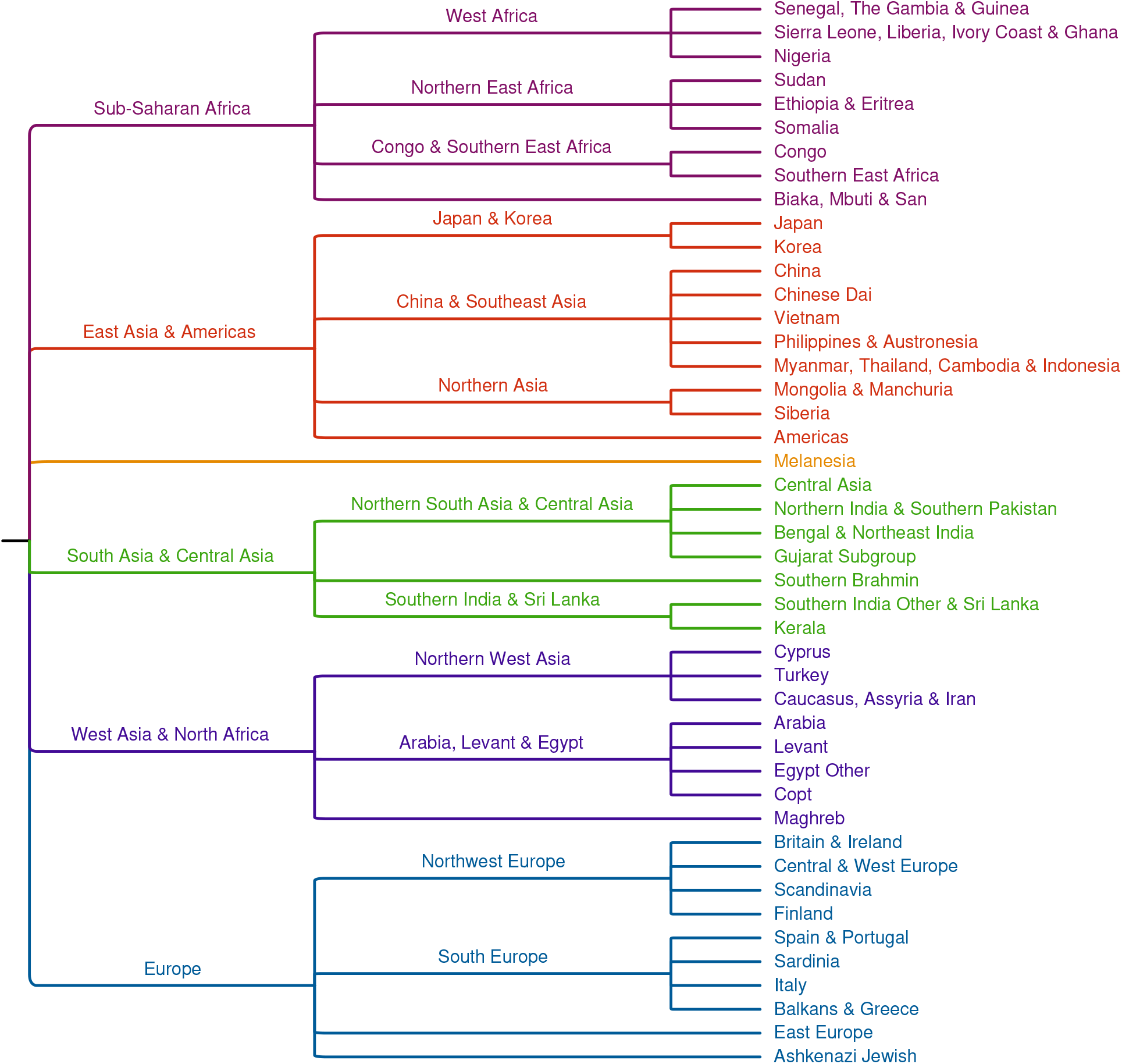
Population hierarchy with 45 reference populations (leaves). Colors reflect the six continental groupings at the highest level of the hierarchy.

For a given haplotype window, we sum the posterior probabilities from the leaves to the root of the tree, so that each node is assigned a probability equal to the sum of its children’s probabilities, with the probability at the root always equal to one. We assign each haplotype window to the lowest node (i.e., the node closest to the leaves) at which the posterior probability exceeds a specified precision level, *t* ∈ [0.5, 1). In the worst case, no node other than the root has a posterior probability exceeding *t*. In this case, we do not classify the window. If assignment probabilities are well calibrated, this procedure ensures that the precision of the assignment is at least *t*. Therefore, we refer to *t* as the “nominal precision threshold”.

### 2.4 Parameter estimation and model evaluation

We used a stratified five-fold cross-validation approach to estimate parameters and evaluate models, maintaining similar representation among the 45 reference populations within each fold. For each fold, we estimated local-classifier parameters using a training set composed of the ~80% of individuals assigned to the other folds. We then classified each window of each chromosome copy of each individual using the models trained with the individual held out, yielding hard-clustering vectors. To estimate emission parameters, including the autoregression transition matrix *ε*, we use a modified supervised EM algorithm applied to these hard-clustering vectors.

In the original implementation of Ancestry Composition (Durand et al., 2014), we estimated APHMM transition parameters using an unsupervised EM training procedure that relied on the natural admixture found in a broader set of 23andMe customers. Specifically, we estimated transition parameters for samples of ~1000 unrelated 23andMe customers from each of the following population groups: African-American, Ashkenazi Jewish, East Asian, European, Latino, Middle-Eastern, and South Asian. We term these groups “smoother training pools”. For each individual to whom we applied Ancestry Composition prediction, we combined the predictions of each smoother training pool’s model using Bayesian model averaging.

In the updated version of our algorithm, we estimate distinct APHMM transition parameter values for each individual to whom we apply Ancestry Composition prediction. To do so, we use the same EM algorithm that was used to estimate transition parameters for the smoother training pools, but, rather than aggregating expectations across ~2000 haplotypes for each chromosome, we aggregate across each individual’s 23 pairs of hard-clustering vectors. To encourage convergence to sensible transition parameter values, we initialize transition-parameter optimization from the pretrained transition parameter sets of the smoother training pools. To determine which smoother training pool is to provide the initial values for transition parameter optimization, we use a multinomial Naive Bayes classifier trained on the hard-clustering assignments of all individuals in all smoother training pools. For each query individual to whom we apply Ancestry Composition, we initialize transition parameter values with those of the smoother training pool chosen by the Naive Bayes classifier when applied to the query individual’s hard-clustering vectors. We find that this individualized transition-parameter optimization affords the error-correction module a great degree of flexibility and, in so doing, reduces bias and increases accuracy.

## 3 Results

We evaluated Ancestry Composition’s classification performance using precision and recall measures computed via a five-fold stratified cross-validation experiment (see subsection 2.4). We estimated precision for population *k* as the proportion of windows predicted to derive from population *k* that actually do derive from population *k*, and we estimated recall for population *k* as the proportion of windows truly deriving from population *k* that were predicted to derive from population *k*.

Table 2 shows accuracy results at continental, regional, and sub-regional scales, as described in subsection 2.3. At each level of the population hierarchy, we estimated precision and recall for two precision thresholds: *t* ∈ {0.5, 0.8}. Note that increasing the threshold increases precision at the expense of recall.

**Table 2:** Precision and recall (%) for all populations in the population hierarchy.

| | Precision<br>$t = 0.5$ | Recall<br>$t = 0.5$ | Precision<br>$t = 0.8$ | Recall<br>$t = 0.8$ |
| --- | --- | --- | --- | --- |
| Sub-Saharan Africa | 99.0 | 98.6 | 99.2 | 98.2 |
| West Africa | 98.4 | 98.9 | 99.0 | 98.1 |
| Senegal, The Gambia & Guinea | 94.5 | 96.0 | 97.0 | 91.3 |
| Sierra Leone, Liberia, Ivory Coast & Ghana | 96.6 | 87.4 | 98.6 | 74.8 |
| Nigeria | 91.6 | 98.9 | 96.7 | 95.3 |
| Northern East Africa | 97.5 | 92.7 | 98.2 | 91.1 |
| Sudan | 95.0 | 83.8 | 96.3 | 79.9 |
| Ethiopia & Eritrea | 93.9 | 97.8 | 95.8 | 97.1 |
| Somalia | 99.0 | 91.6 | 99.3 | 90.4 |
| Congo & Southern East Africa | 97.4 | 99.4 | 98.5 | 98.2 |
| Congo | 98.1 | 99.3 | 99.2 | 97.6 |
| Southern East Africa | 93.5 | 97.1 | 96.6 | 92.7 |
| Biaka, Mbuti & San | 98.5 | 86.3 | 99.2 | 80.2 |
| East Asia & Americas | 98.7 | 99.5 | 99.0 | 99.3 |
| Japan & Korea | 98.6 | 99.9 | 99.0 | 99.9 |
| Japan | 99.7 | 99.2 | 99.7 | 99.0 |
| Korea | 96.0 | 99.6 | 97.0 | 99.5 |
| China & Southeast Asia | 99.6 | 98.0 | 99.7 | 97.0 |
| China | 96.7 | 94.3 | 97.5 | 91.4 |
| Chinese Dai | 81.6 | 98.2 | 87.4 | 97.3 |
| Vietnam | 94.4 | 98.0 | 96.7 | 97.2 |
| Philippines & Austronesia | 93.8 | 89.6 | 94.7 | 86.5 |
| Myanmar, Thailand, Cambodia & Indonesia | 94.9 | 66.5 | 95.8 | 57.9 |
| Northern Asia | 56.0 | 95.0 | 67.3 | 92.3 |
| Mongolia & Manchuria | 38.1 | 91.6 | 50.7 | 86.1 |
| Siberia | 91.9 | 95.1 | 95.3 | 92.4 |
| Americas | 98.8 | 94.1 | 99.3 | 90.3 |
| Melanesia | 98.8 | 98.3 | 99.2 | 96.9 |
| South Asia & Central Asia | 98.5 | 96.9 | 98.9 | 95.9 |
| Northern South Asia & Central Asia | 94.3 | 91.6 | 95.8 | 88.7 |
| Central Asia | 86.1 | 49.4 | 88.5 | 37.4 |
| Northern India & Southern Pakistan | 82.3 | 87.8 | 85.5 | 83.0 |
| Bengal & Northeast India | 91.6 | 94.7 | 94.6 | 92.6 |
| Gujarat Subgroup | 98.2 | 97.7 | 98.7 | 96.8 |
| Southern Brahmin | 93.3 | 83.9 | 95.0 | 80.1 |
| Southern India & Sri Lanka | 89.8 | 94.2 | 92.3 | 91.9 |
| Southern India Other & Sri Lanka | 78.1 | 93.3 | 82.1 | 90.6 |
| Kerala | 92.8 | 75.5 | 94.6 | 73.5 |
| West Asia & North Africa | 95.8 | 95.3 | 97.1 | 93.4 |
| Northern West Asia | 83.7 | 92.0 | 87.7 | 89.0 |
| Cyprus | 94.7 | 93.3 | 97.0 | 90.2 |
| Turkey | 86.2 | 67.6 | 91.8 | 56.2 |
| Caucasus, Assyria & Iran | 67.1 | 92.9 | 74.6 | 88.8 |
| Arabia, Levant & Egypt | 94.3 | 85.6 | 95.8 | 81.8 |
| Arabia | 88.2 | 70.3 | 91.2 | 66.1 |
| Levant | 93.8 | 70.5 | 95.7 | 63.4 |
| Egypt Other | 74.1 | 90.2 | 82.1 | 86.4 |
| Copt | 92.4 | 95.3 | 95.1 | 93.7 |
| Maghreb | 97.4 | 89.9 | 98.4 | 86.7 |
| Europe | 98.7 | 99.0 | 99.0 | 98.4 |
| Northwest Europe | 96.1 | 96.3 | 97.2 | 94.0 |
| Britain & Ireland | 95.4 | 88.5 | 98.2 | 75.4 |
| Central & West Europe | 79.7 | 81.3 | 87.1 | 64.4 |
| Scandinavia | 90.8 | 90.7 | 94.9 | 81.9 |
| Finland | 93.5 | 95.3 | 95.3 | 93.3 |
| South Europe | 91.1 | 89.7 | 94.0 | 85.2 |
| Spain & Portugal | 90.0 | 96.5 | 94.1 | 94.0 |
| Sardinia | 88.1 | 95.2 | 92.3 | 92.3 |
| Italy | 87.8 | 86.1 | 92.1 | 79.1 |
| Balkans & Greece | 89.5 | 81.0 | 92.5 | 75.0 |
| East Europe | 85.9 | 88.8 | 89.4 | 82.6 |
| Ashkenazi Jewish | 99.3 | 98.6 | 99.4 | 98.1 |

At the continental scale (i.e., for all non-leaf populations that are children of the root node), when *t* = 0.5, precision exceeds 97% and recall exceeds 92%. When *t* = 0.8, precision is greater than 99% for all continents except Europe, which achieves a precision of 98.3%, and recall drops slightly, with a minimum of 89.4% for West Asia and North Africa.

At the regional scale (i.e., considering the twelve non-continent non-leaf populations), precision and recall are uniformly less than or equal to the continent-scale parent populations, by definition. With nominal precision threshold *t* = 0.5, precision remains fairly high, with median of 95.2%, and it exceeds 90% for all but North Asia and Northwest Asia. Recall for *t* = 0.5 is also relatively good at this scale, with median of 94.5%. It exceeds 90% for eight of twelve regions and exceeds 80% for all. At nominal precision threshold *t* = 0.8, precision has a median of 97.5% and is greater than 90% for all regions except North Asia and Northwest Asia. Recall decreases slightly but still remains above 85% for most populations.

At the leaf level, many populations continue to have good precision and recall metrics. Seven of 45 leaf populations (namely, Ethiopia & Eritrea, Congo, Japan, Korea, China, Gujarati Subgroup, and Ashkenazi) achieve precision and recall greater than 95% for both nominal precision thresholds, and 17 of 45 leaf populations achieve precision and recall greater than 90% for both precision thresholds. At nominal precision threshold *t* = 0.5, the median precision is 96.1% for all leaf-level populations, with 35/45 leaf-level populations having precision at least 90% and 41/45 populations having precision at least 80%. Recall at *t* = 0.5 has median 91.1%, with 26/45 leaf-level populations exceeding 90% and 38/45 populations exceeding 80%. As at the continental and regional scales, precision increases slightly and recall decreases slightly with nominal precision threshold *t* = 0.8, as compared to *t* = 0.5.

## 4 Discussion

We have developed a two-stage pipeline for ancestry deconvolution, Ancestry Composition. This modular approach makes our method flexible, robust, and easy to update. Ancestry Composition achieves high precision for closely related populations, as demonstrated by our cross-validation experiments, and it outputs probabilistic assignments, which enable confidence-threshold tuning and hierarchical classification. Unlike many previous approaches that can only distinguish between a few well-differentiated populations, Ancestry Composition can be trained to differentiate a large number of closely related populations.

The ultimate goal of ancestry deconvolution is to estimate *P* (*Y* | *X*), the distribution of unobserved ancestry states *Y*, given observed haplotypes *X*. Generative approaches, such as HMMs, achieve this by first estimating the joint distribution *P* (*X, Y*) and then conditioning on the observed data, *X*. In contrast, discriminative approaches directly model the conditional distribution *P* (*Y* |*X*).

Discriminative approaches to classification often outperform generative methods (Lafferty et al., 2001), and the additional complexity required to fully model *P* (*X, Y*) tends to limit generative ancestry deconvolution methods to just a few ancestral populations. In addition, discriminative approaches are typically more robust to model misspecification, precisely because they do not attempt to fully model the joint distribution of haplotypes and their ancestries. In light of these advantages, our local classifier does not assume any particular demographic model underlying the admixture process. Rather, our SKSVM learns decision boundaries between the reference populations directly from the data.

Despite the advantages discriminative approaches offer, generative models are generally more flexible and permit expression of more complex dependencies between observations and hidden random variables. Therefore, Ancestry Composition adopts a mixed approach, as advocated in Jaakkola et al. (1999) and Ng and Jordan (2002), in which the output of a discriminative local classifier is input to a generative error-correction module implemented as an autoregressive pair hidden Markov model. The hidden-Markov-model framework provides a natural means by which to correct phasing switch errors and model dependencies between adjacent observations.

In contrast to our mixed approach, RFMix (Maples et al., 2013) is a purely discriminative approach for admixture deconvolution. It implements random forests as its local classifier and conditional random fields to reconcile adjacent chromosomal windows. Random forests have some advantage over SVMs; they are inherently multiclass classifiers, and they output probabilities rather than hardcalls. However, SVMs offer a direct way to plug in a kernel, which enables us to efficiently extract an enormous number of features from short chromosomal segments.

Other purely discriminative approaches have recently been developed. Kumar et al. (2020) have described an approach that employs boosted gradient trees to perform local ancestry inference much faster and with fewer computational resources than existing methods, while maintaining comparable accuracy. A similar method, using neural networks has also been described recently (Montserrat et al., 2020).

## 5 Acknowledgments

We thank Brian Naughton, Katarzyna Bryc, Nicholas Eriksson, Robert Bell, Ethan Jewett, Will Freyman, Kimberly McManus, and other members of the 23andMe Ancestry R&D team for insightful comments throughout the development of this project. We also thank the many 23andMe customers who answered survey questions and allowed us to study their genomes; this work would not have been possible without them.

